# Progressive Loss of Astrocytic AIBP Expression during Alzheimer’s Disease Pathology

**DOI:** 10.1101/2025.10.03.680289

**Authors:** Shihui Ding, Soo-Ho Choi, Yi Sak Kim, Nicolaus Nazarenkov, Yury I. Miller

## Abstract

Astrocytes and microglia play crucial roles in mediating neuroinflammation during Alzheimer’s disease (AD) progression. ApoA-I binding protein (APOA1BP, also known as AIBP/NAXE) attenuates neuroinflammation by blocking amyloid β-induced TLR4 inflammaraft formation and oxidative stress. *Apoa1bp* knockout in APP/PS1 mice exacerbates microgliosis, increases amyloid plaque burden, neuronal cell loss, and reduces survival at 6 months. Although *APOA1BP* mRNA is ubiquitously expressed in humans, its cell-type-specific distribution in the brain remains unclear. To examine AIBP protein expression in the human brain, we performed immunohistochemistry on hippocampal sections from postmortem brain specimens from subjects aged 75-96 of both sexes. Using GFAP and IBA1 to label astrocytes and microglia, respectively, we found that AIBP protein was highly expressed in astrocytes, but not in microglia. Stratification of subjects by Braak stage (I-II, III-IV, V-VI) revealed a progressive decline in astrocytic AIBP expression with advancing AD pathology. Meta-analysis of RNA-seq profiling indicated enriched *Apoa1bp* expression in adult mouse astrocytes. Systemic *Apoa1bp* knockout in the APP/PS1 mouse exacerbated astrogliosis. These findings demonstrate that AIBP is predominantly expressed in astrocytes and its expression declines with AD progression, suggesting a potential role for AIBP in astrocyte-mediated neuroprotection and AD pathogenesis.

## Introduction

Glial cells, including microglia and astrocytes, orchestrate neuroinflammatory responses in Alzheimer’s disease (AD). While microglia act as the resident immune cells of the brain and contribute to amyloid beta (Aβ) clearance and inflammatory signaling, astrocytes, the most abundant cell type in the central nervous system, have emerged as key modulators of excitotoxicity, oxidative stress, lipid metabolism and neuroinflammation. Current studies suggest that astrocytes exert both protective and detrimental effects depending on disease stage and microenvironmental factors. Therapeutic strategies targeting astrocytes have therefore focused on modulating anti-inflammatory and antioxidant pathways, regulating glutamate activity, and restoring lipid and calcium homeostasis [1, 2].

ApoA-I binding protein (APOA1BP, also known as AIBP or NAXE) is a protein originally identified in a screen for apoA-I interacting proteins [3]. AIBP regulates cholesterol efflux and the molecular architecture of inflammarafts, defined as cholesterol-rich, enlarged lipid rafts chronically expressing assembled complexes of inflammatory receptors, including TLR4 dimers [4]. Experimental evidence demonstrates that AIBP mitigates neuroinflammation by protecting against Aβ-induced TLR4 inflammaraft formation and oxidative stress [5]. In *Apoa1bp*-deficient APP/PS1 mice, microgliosis is exacerbated, leading to increased Aβ plaque burden, neuronal cell death, and reduced survival at 6 months of age [5]. These findings position AIBP as a candidate therapeutic for mitigating Aβ-induced TLR4-dependent inflammatory responses as AD advances.

Most evidence for AIBP’s potential protective effects in AD arises from cell biology and mouse studies [5, 6]; whether the same mechanisms operate in human tissue remains unknown. At the cellular level, AIBP can be both secreted or localized to the mitochondria. Transcriptomic data indicate that human AIBP mRNA is broadly expressed across tissues, with particularly high levels in kidney, liver, brain, testis, thyroid, and adrenal gland [7]. However, the cell-type specificity of AIBP expression in the human brain is poorly characterized. Given that the hippocampus—a region critical for learning and memory—is among the earliest affected brain regions in AD [8], here we investigated AIBP expression across hippocampal cell types to determine whether cell-type specific expression is associated with AD pathology.

## Methods

### Human Tissue Specimens

Formalin-fixed paraffin embedded postmortem human brain sections across various Braak stages were obtained from the brain bank of the University of California, San Diego Shiley-Marcos Alzheimer’s Disease Research Center (UCSD ADRC). Hippocampus sections of 14 postmortem brain specimens from subjects aged 75–96 of both sexes were used.

### Immunohistochemisty

Formalin-fixed paraffin embedded human brain sections were deparafinized in Histoclear three times for 5 min each and rehydrated for 3 min in a graded series of ethanol concentrations (100%, 100%, 90%, 80%, 70%, 50%, and 10%). For antigen retrieval, slices were heated in a preheated water bath with 10 mM sodium citrate (pH 6.0) at 95 °C for 30 min. The container was then cooled for 20 min at room temperature (RT). Subsequently, the slices were washed three times in Tris-buffered saline with 0.05% Tween-20 (TBS-T) for 5 min each and blocked in a protein block (ab64226; Abcam) at RT for 1 h. The antibodies used were anti-AIBP (20 μg/ml, 1:190, BE-1, produced in-house [9-11]), anti-GFAP (1:200, 53-9892-82; Invitrogen), and anti-IBA1 (1:500, 013-26471; Fujifilm Wako). The slices were incubated with the anti-AIBP antibody overnight at 4 °C and washed five times in TBS-T for 5 min each. Secondary antibody (1:1000, A-11004; Invitrogen) was applied for approximately 2 h at RT. After washing with TBS-T, the slices were incubated with anti-GFAP and anti-IBA1 antibodies overnight at 4 °C. Slices were then incubated with DAPI and washed five times in total. To reduce nonspecific background staining, slides were rinsed with TBS without detergent four times and then incubated in a TrueBlack Lipofuscin Autofluorescence Quencher (23007; Biotium) for 30 s. Slices were finally washed three times with TBS, mounted with ProLong Glass Antifade Mountant (P36984; Invitrogen) and coverslipped.

### Mouse brain immunohistochemistry

Mouse experiments were conducted in accordance with the protocol approved by the Institutional Animal Care and Use Committee of the University of California, San Diego. *Apoa1bp*^*-/-*^ APP/PS1 mice were created in house as we reported [5]. Housing conditions, tissue preparation and immunohistochemisty followed the procedures described in our recent paper [5]. An anti-Aβ antibody (1:100, 82E1, IBL 10323) and an anti-GFAP antibody (1:200, 53-9892-82; Invitrogen) were used for immunohistochemisty.

### Imaging analysis

Fluorescent images of stained human brain sections were acquired using a Leica SP8 confocal microscope using a ×20/0.75 NA dry objective with a 2048 × 2048 pixel field of view. Images of mouse brains were acquired using a Leica SP8 confocal microscope operated in a Lightning deconvolution mode. Colocalization analysis was performed with ImageJ using the Coloc 2 plugin. Pearson correlation coefficient (*r*) and Manders’ coefficients (above zero intensity of Channel 2) were generated for the region of interest within the same field of view (Fig. 1). Image analysis was performed using ImageJ. Cell boundaries were segmented by applying the Moments threshold and the Analyze Particles function in the GFAP channel. The mean intensity of each cell was calculated and plotted as histogram (Fig. 2B). A threshold of 1.5 a.u. for mean intensity was used to identity AIBP-positive astrocytes in Fig. 2C. All image processing and quantification were carried out in a blinded manner. All statistical analyses were conducted using GraphPad Prism, version 10.

**Figure 1.**
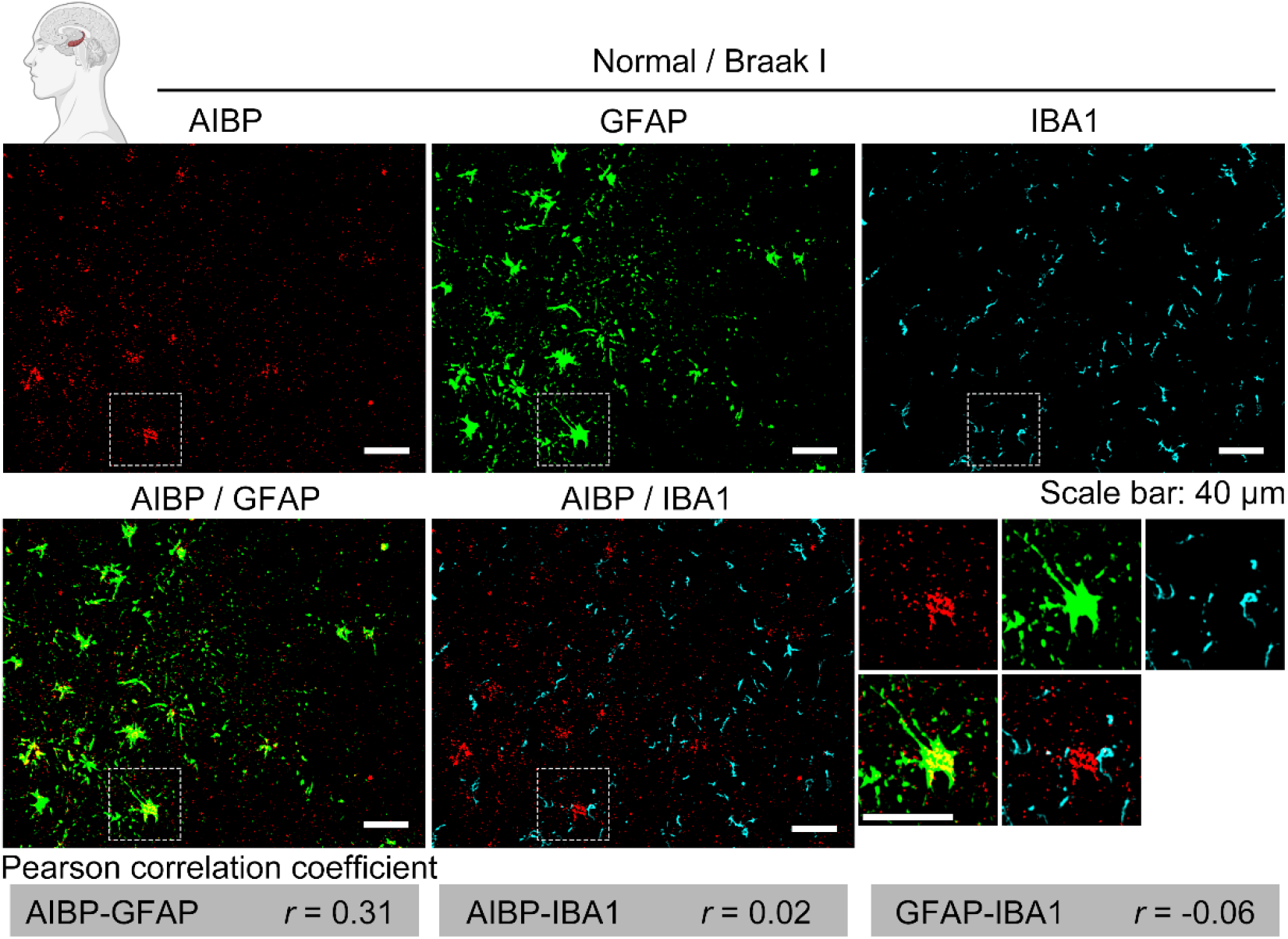
AIBP protein is highly expressed in astrocytes in hippocampal tissue from humans. Representative images showing AIBP (red) expression in GFAP-labeled astrocytes (green) and IBA1-labeled microglia (cyan) in paraffin-embedded normal human brain with Braak stage I. White dashed boxes indicate regions shown at higher magnification. Pearson correlation coefficient (*r*) was calculated between each pair of channels. Scale bar, 40 μm.

**Figure 2.**
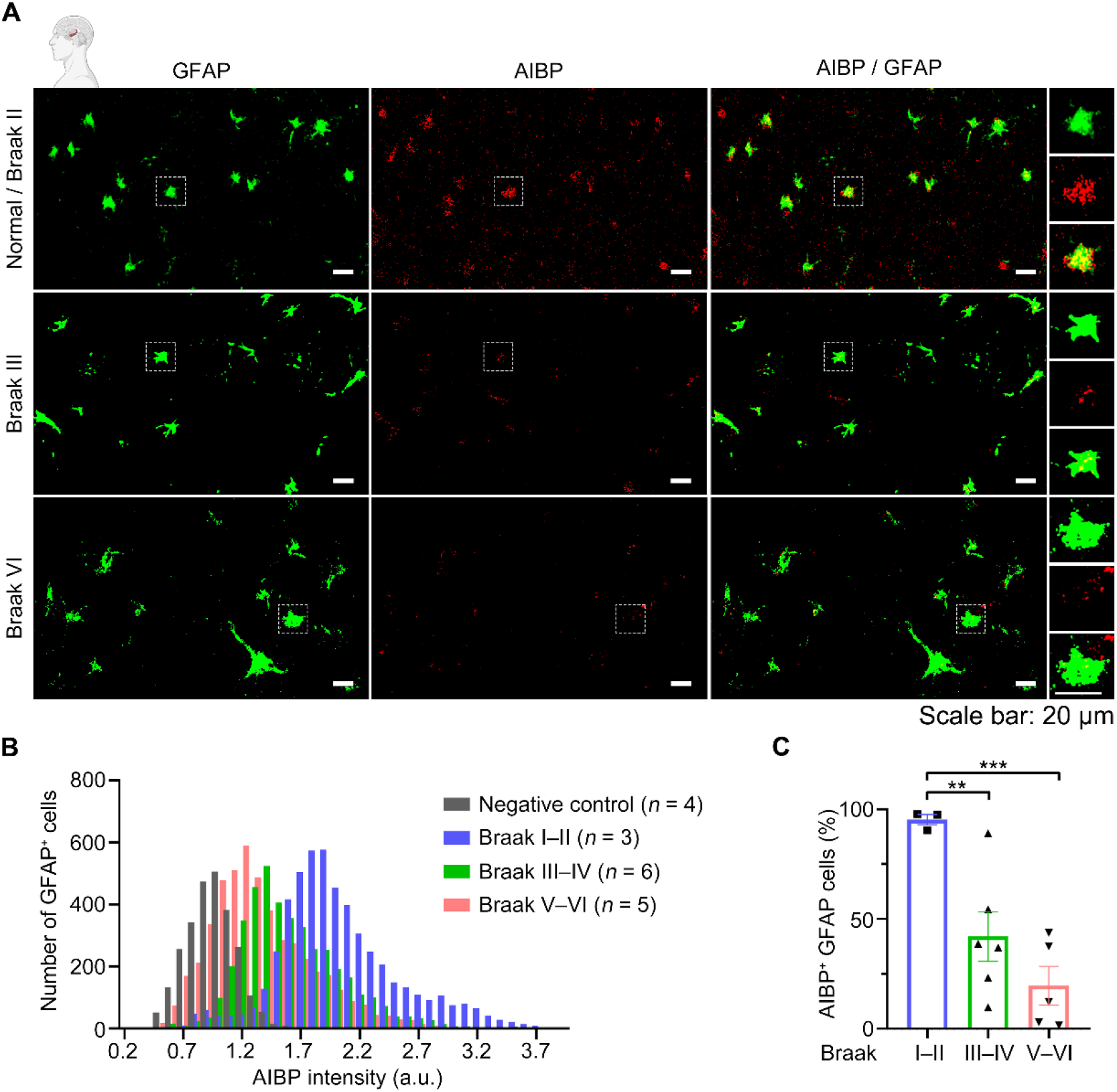
Astrocytic AIBP expression shows a progressive decline with advancing AD pathology. **A**, Representative images showing AIBP (red) expression in GFAP-labeled astrocytes (green) in the hippocampus of human brains across different Braak stages. White dashed boxes indicate regions shown at higher magnification. Scale bar, 20 μm. **B**, Histogram of AIBP expression levels in hippocampal GFAP^+^ astrocytes across different Braak stages. Negative control lacks the AIBP primary antibody, with other conditions remaining unchanged. Braak I–II, normal or individuals without pathology. Braak III–IV, patients with dementia. Braak V–VI, AD patients. *n*, patients number in each group. **C**, Percentage of AIBP^+^ cells in GFAP^+^ astrocytes with AIBP intensity thresholds set at > 1.5 a.u. Each dot represents an individual patient. Mean ± SEM. **, p<0.01. ***, p<0.001. Patients include both genders.

## Results

### AIBP is predominantly expressed in astrocytes in the human hippocampus

To identify the cell-type sepecfic AIBP expression, we performed immunohistochemistry on hippocampal sections from postmortem human brains without AD pathology, using GFAP and IBA1 to label astrocytes and microglia, respectively. We found that AIBP signal was highly colocalized with GFAP (**Fig. 1**; Pearson’s correlation coefficient, *r* = 0.31). In contrast, little to no colocalization was observed between AIBP and IBA1 (*r* = 0.02). The correlation between GFAP and IBA1 was −0.06, confirming the specificity of the two glial markers. Simiarly, Manders’ M1 coefficients for AIBP–GFAP, AIBP–IBA1, and GFAP–IBA1 were 0.602, 0.210, and 0.139, respectively. Although no neuronal countersatining was performed, the pattern of AIBP staining suggested no detectable neuronal AIBP expression under the conditions of this experiment. No signal was detected in the secondary antibody-only negative control omitting the AIBP primary antibody (**Fig. S1**). Together, these results demonstrate the predominant expression of AIBP in astrocytes within the hippocampus of the human brain.

### Astrocytic AIBP expression shows a progressive decline with advancing AD pathology

Reactive astrogliosis is a hallmark of Alzheimer’s disease and is characterized by profound transcriptional, morphological, and functional changes in astrocytes [12]. Given that astrocytes express high levels of AIBP (**Fig. 1**), we asked whether AD pathology affects astrocytic AIBP expression in human brains. To address this, we performed immunohistochemistry on hippocampal sections from 14 postmortem human brains (aged 75–96 years, both sexes). The subjects were grouped by Braak stages (I–II, III–IV, V–VI).

When comparing astrocytic AIBP expression across Braak stages, we observed a progressive decline as AD pathology advanced (**Fig. 2A**). Astrocytic AIBP expression was highest at Braak I–II, significantly reduced at Braak III–IV, and reached the lowest levels at Braak V–VI (**Fig. 2B**). When applying a threshold to define AIBP-positive astrocytes, their number was significantly reduced in Braak V–VI compared to Braak I–II, with an intermediate reduction at Braak III–IV (**Fig. 2C**).

Microglial activation is another characteristic feature of AD pathology, characterized by the transformation of ramified homeostatic microglia into amoeboid, activated microglia. Accordingly, we examined AIBP expression in relation to IBA1-labeled microglial morphology across Braak stages (**Fig. 3**). Compared with Braak II, Braak VI samples showed an overall decline in AIBP expression, despite limited colocalization with IBA1-positive microglia, accompanied by less ramified and more amoeboid microglial morphology.

**Figure 3.**
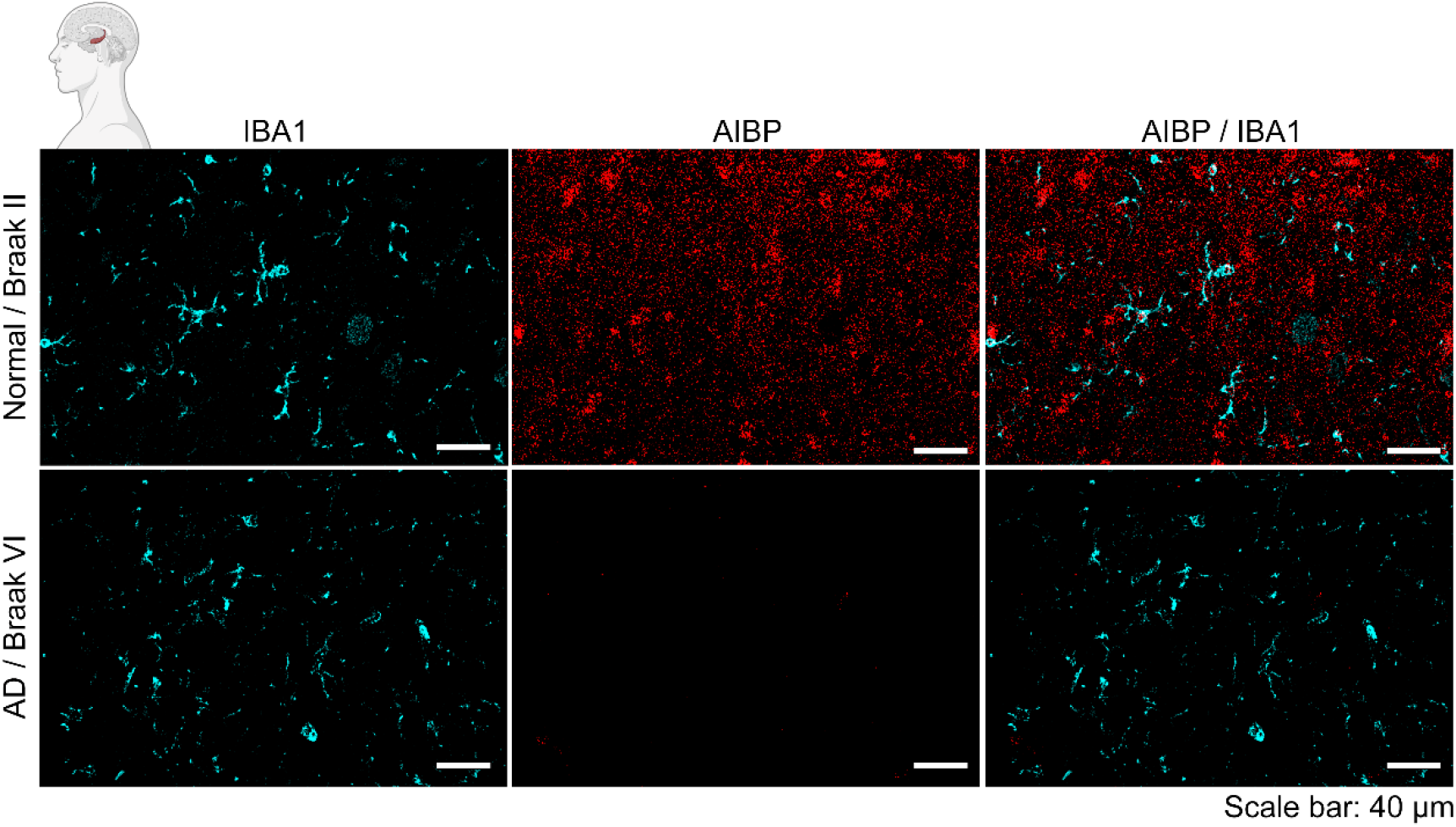
AIBP expression and microglia morphology in AD pathology. Representative images showing AIBP (red) expression and IBA1-labeled microglia (cyan) in the hippocampus of human brains at different Braak stages. Braak II, normal or individuals without pathology. Braak IV, AD patients. Scale bar, 40 μm.

Taken together, these findings demonstrate that AIBP expression progressively declines as AD pathology advances, highlighting a potential role for AIBP in astrocyte function and its relevance to AD pathogenesis.

### *Apoa1bp* is highly expressed in adult mouse astrocytes, and AIBP deficiency augments astrogliosis in the APP/PS1 AD mouse model

To understand astrocytic AIBP expression pattern in a mouse, we searched for the mouse expression profile of *Apoa1bp* using adult astrocytic RNA-seq explorer [13, 14]. We found that *Apoa1bp* gene was predominantly expressed in astrocytes relative to whole tissue across the hippocampus, striatum and cortex (**Fig. 4**). We also observed a simlar enriched expression of *Gfap* and *Abca1*, while *Cx3cr1, Tmem119 and Map2* showed the opposite trend (not shown), suggesting that the RNA profile contains minimal contamination from microglia and neurons. Techical issues of using a mouse monoclonal anti-AIBP antibody did not allow us to examine AIBP protein expression in the mouse brain.

**Figure 4.**
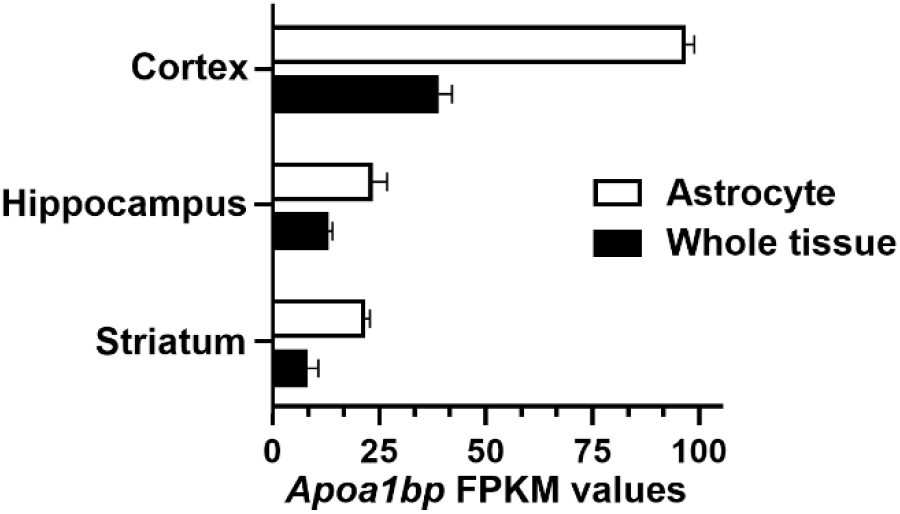
*Apoa1bp* expression in adult astrocyte. Meta-analysis of region-specific astrocytic RNA-seq from 8–12-week-old adult mice (2 males and 2 females) [13, 14] revealed that astrocytic *Apoa1bp* expression was predominant across several brain regions including cortex, hippocampus, and striatum, compared with whole-tissue expression. FPKM, Fragments Per Kilobase of transcript per million mapped reads. Mean + SEM.

Importantly, when we knocked out the *Apoa1bp* gene in the APP/PS1 AD mouse model [5, 15], astrocyte density in the hippocampus increased along with elevated Aβ signals (**Fig. 5A and 5B**). These results indicate that AIBP deficiency in the APP/PS1 model exacerbates astrogliosis and aggravates AD pathology.

**Figure 5.**
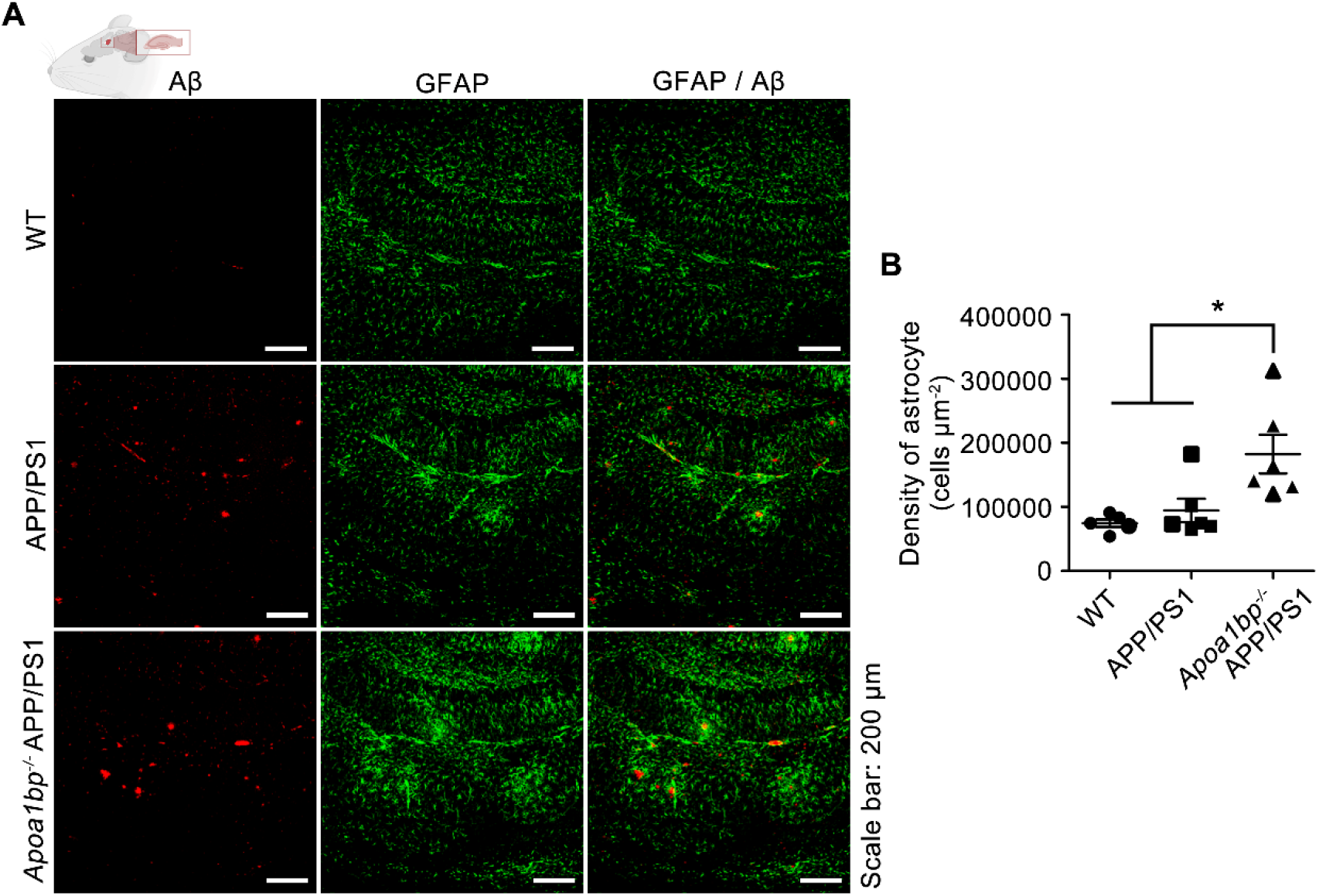
Astrocyte density increased in the hippocampus of *Apoa1bp*^*-/-*^ mice on the APP/PS1 background. **A**, Representative images showing Aβ plaque (red) and GFAP-labeled astrocytes (green) in the hippocampus of mouse brains in WT, APP/PS1, and *Apoa1bp*^*-/-*^ APP/PS1 mice. Scale bar, 200 μm. **B**, Statistical analysis of GFAP-labeled astrocyte density in the hippocampus of WT (*n* = 5), APP/PS1 (*n* = 6), *Apoa1bp*^*-/-*^ APP/PS1 (*n* = 6) mice. Mean ± SEM. *, p<0.05. One-way ANOVA with Bonferroni’s multiple comparison test.

## Discussion

The AIBP expression varies under different pathological conditions. It is increased in atherosclerotic lesions and in oxidized LDL-stimulated macrophages [9], as well as in inflammatory cells in the human lung, and is secreted into the bronchoalveolar space following LPS inhalation in mice [16]. However, AIBP is downregulated in asthmatic bronchial epithelial cells [11] and in the glaucomatous retina [17]. Given the distinct roles of neurons, astrocytes, microglia, and oligodendrocytes in neurological maintenance and cognitive resilience, it remained unclear whether AIBP expression is uniform or varies in a cell-type-specific manner. Defining AIBP cellular distribution in human brain tissue is therefore important to link physiological effects with cell-specific functions and assess its role in Alzheimer’s disease.

In this study, we mapped AIBP expression in astrocytes and microglia in the hippocampus of human brains. We found that the AIBP protein was highly expressed in astrocytes and to a lesser degree in microglia. Importantly, astrocytic AIBP expression progressively declined with advancing AD pathology, suggesting a protective role for AIBP in astrocyte function. In agreement, a meta-analysis using the mouse brain RNA-seq explorer [13, 14] showed the predominant *Apoa1bp* mRNA expression in astrocytes across the hippocampus, striatum and cortex.

Consistently, a meta-analysis of single-nucleus RNA-seq [18] from the prefrontal cortex of 48 individuals with varying degrees of AD pathology showed that neuronal *APOA1BP* mRNA expression also declined progressively with disease severity (**Fig. 6**). The reduction was most pronounced in late-stage AD, with intermediate decreases in early-stage AD, and occurred in both excitatory and inhibitory neurons. Thus, AIBP reduction during AD progression is observed at the mRNA and protein levels, in neurons and astrocytes, further supporting the potential protective role of AIBP in the human brain.

**Figure 6.**
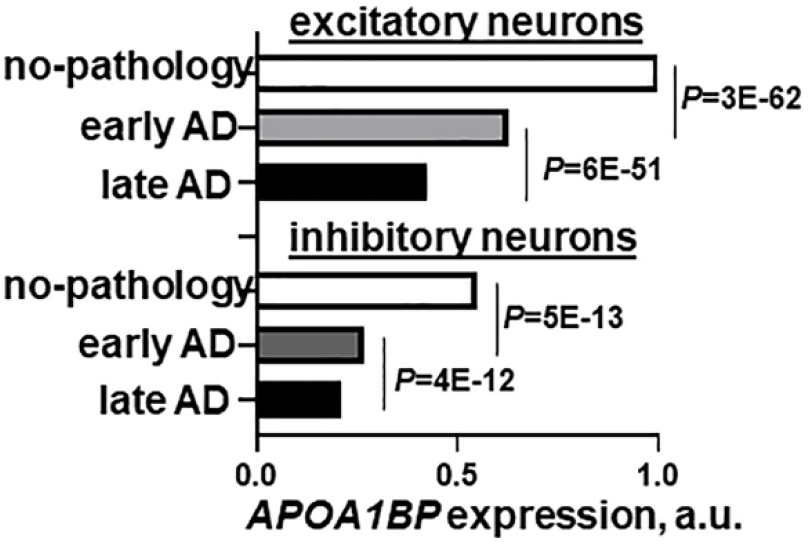
AIBP expression in human AD brain. Meta-analysis of single-nucleus RNA-seq [18] showed *APOA1BP* expression is reduced in early and late AD.

To investigate the role of AIBP in the brain, we deleted the *Apoa1bp* gene in the APP/PS1 AD mouse model. Our previous study demonstrated that *Apoa1bp*^*-/-*^ APP/PS1 mice developed more amyloid plaques and exhibited enhanced microgliosis compared with APP/PS1 mice [5]. In the present study, we further observed exacerbated astrogliosis in *Apoa1bp*^*-/-*^ APP/PS1 mice. A drawback of this model is that *Apoa1bp* deficiency was global rather than astrocyte-specific, raising the possibility that loss of *Apoa1bp* in other cell types also contributed to the aggravated AD pathology. Although the link between predominant *Apoa1bp* expression in astrocytes and the observed astrogliosis remains unclear, our findings suggest that *Apoa1bp* deficiency induces astrocytic dysfunction in the context of AD. Future studies employing astrocytic-specific genetic manipulation, such as conditional knockout or targeted replenishment of AIBP in APP/PS1 mice, would provide more definitive insights.

The limitations of this study also include a limited number of human brain specimens examined and apparently a low sensitivity of the AIBP antibody detection, given that no significant neuronal or microglial expression was detected. However, the finding of a predominant AIBP expression in astrocytes is important for further evaluation of the role a cell-specific AIBP expression plays in neuroprotection during the development of Alzheimer’s disease.

## Author Contributions

Y.I.M. and S.D. conceived the project, Y.I.M., S.D., and S.-H.C. designed the experiments, S.D., Y.S.K. and N.N. performed experiments, S.-H.C. contributed to study discussions. S.D. and Y.I.M. wrote the manuscript.

---

**Figure 1S.**
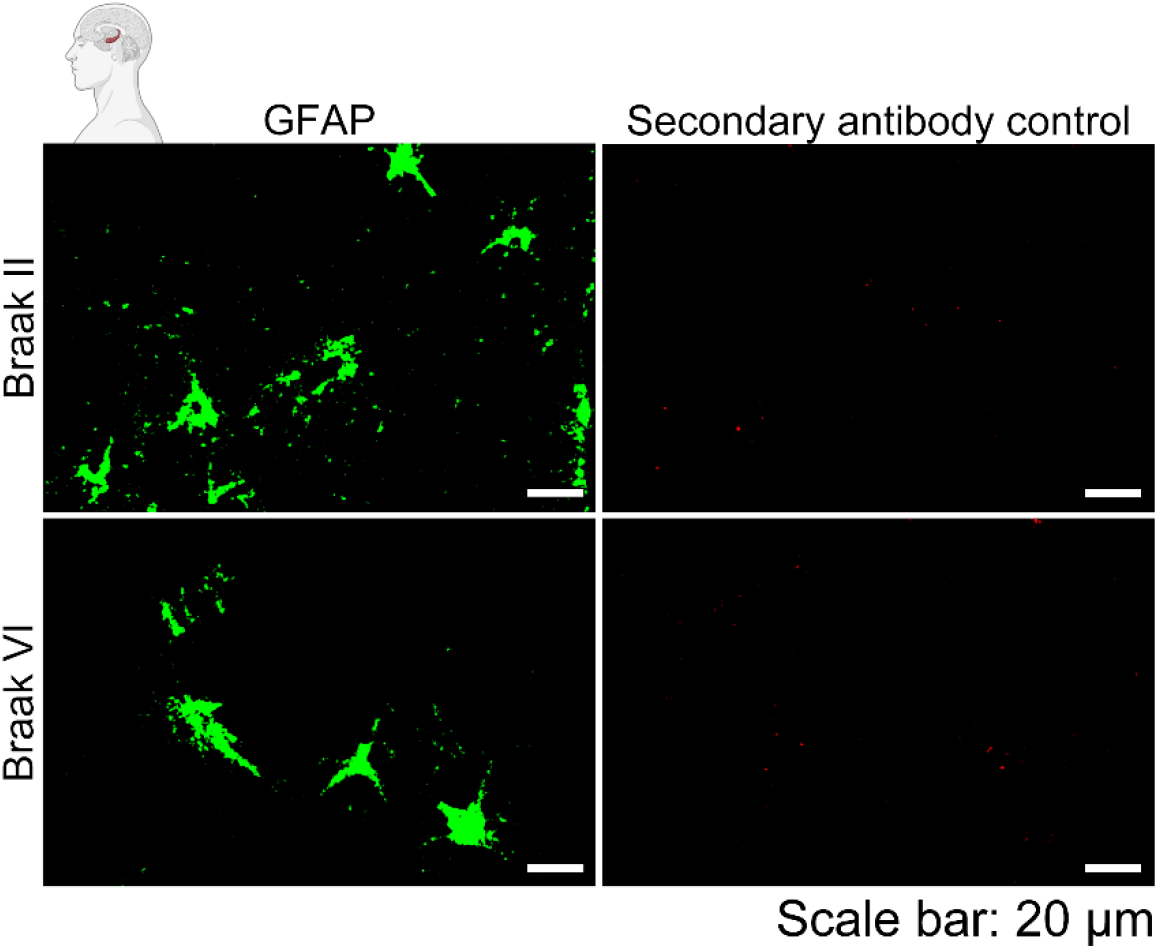
Secondary antibody–only negative control for AIBP immunohistochemistry. Representative images showing secondary antibody signal (red) in GFAP-labeled astrocytes (green) from paraffin-embedded human brain in the hippocampus region at different Braak stages. The controls were performed without the AIBP primary antibody, with other conditions remaining unchanged. Braak II, normal or individuals without pathology. Braak IV, AD patients. Scale bar, 20 μm.

